# Direct simulation of a stochastically driven multi-step birth-death process

**DOI:** 10.1101/2021.01.20.427480

**Authors:** Gennady Gorin, Lior Pachter

**Affiliations:** Division of Chemistry and Chemical Engineering, California Institute of Technology, Pasadena, CA, 91125; Division of Biology and Biological Engineering & Department of Computing and Mathematical Sciences, California Institute of Technology, Pasadena, CA, 91125

## Abstract

The description of transcription as a stochastic process provides a framework for the analysis of intrinsic and extrinsic noise in cells. To better understand the behaviors and possible extensions of existing models, we design an exact stochastic simulation algorithm for a multimolecular transcriptional system with an Ornstein-Uhlenbeck birth rate that is implemented via a special function-based time-stepping algorithm. We demonstrate that its joint copy-number distributions reduce to analytically well-studied cases in several limiting regimes, and suggest avenues for generalizations.

## 2 Background

### 2.1 Markov modeling of transcriptional processes

Recent methods in single-cell transcriptomics have enabled increasingly precise measurements of copies of mRNA molecules in cells [1, 2]. These experimental improvements have dovetailed with theoretical and computational improvements in modeling transcription in cells. The chemical master equation (CME) is the standard modeling framework for discrete-valued processes [3], providing a natural representation of biomolecular counts. CME models that can be used to compute entire discrete distributions [4] have therefore become increasingly relevant for modeling transcription in cells, and are being used for the purpose of statistical inference of underlying biophysical parameters [5].

No canonical choice of model exists, but certain conventions and assumptions are common (See Figure 1). In the CME formalism, a cell is generally represented as a continuous-time Markov chain traversing a discrete state space. The transitions between states are determined by a set of rates. If all rates are time-independent, residence time in each state is described by an exponential random variable parameterized by the sum of efflux rates, while the choice of transition is made based on the respective relative rates. Furthermore, the state space of the Markov chain may be multi-dimensional and a state may therefore correspond to a *vector* of quantities. A common modeling choice uses N_0_ as the domain for dimensions representing molecular species and a finite set of integers as the domain for dimensions representing gene states [6].

**Figure 1:**
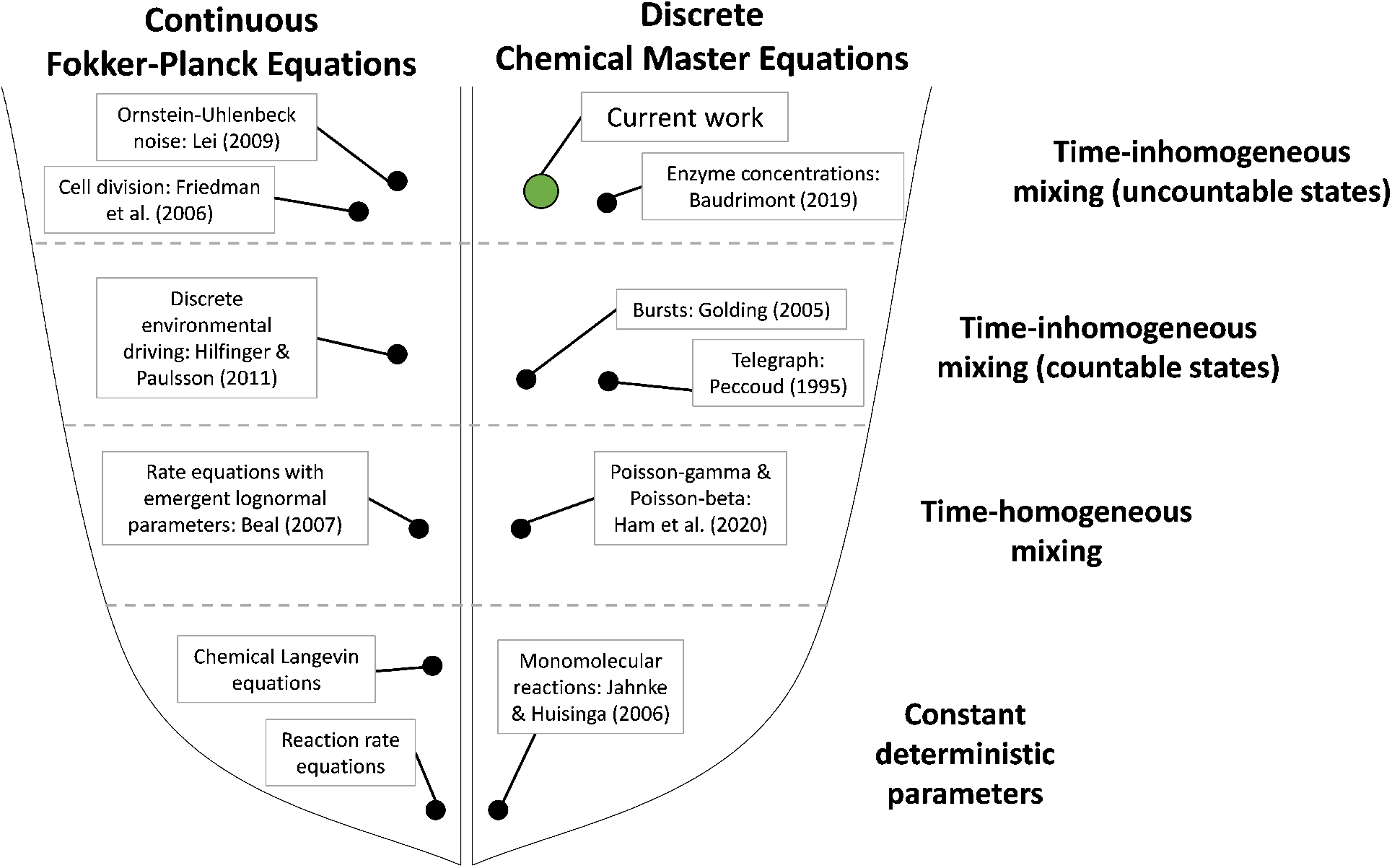
The space of commonly used stochastic models for chemical reactions [2, 6–13].

Given this formalization, it is possible to construct models of gene expression representing physiology of interest. In accordance with recent advances in multimodal data acquisition and modeling [14, 15], we represent mRNA dynamics by a two-stage birth-death process (BDP). A gene locus generates nascent mRNA (unspliced or pre-mRNA) by some variant of a Poisson process. After an exponentially distributed delay with rate *β*, nascent mRNA is converted to mature mRNA (spliced mRNA). Finally, after an exponentially distributed delay with rate *γ*, the mature mRNA is degraded. The choice of a two-stage BDP is informed by the physiological relevance of splicing and buffering models, as well as our recent discussion of qualitative differences in stationary distributions under different noise models [16].

The simplest stochastic description of gene expression, corresponding to unregulated *constitutive*production, has a single gene state [7] that produces mRNA at a constant rate (Figure 2a). Inthe light of activation and deactivation known to occur in many biological systems, a more physiologically relevant model posits two distinct gene states with different mRNA production rates, their switching governed by a telegraph process [6]. A common, and physiologically borne out simplification known as the *bursty approximation* [17] considers the limit of infinitesimally short active periods, which generate finite numbers of gene products [2, 18] (Figure 2b). This description produces statistical over-dispersion over the constitutive production model, an excess of variance that is referred to as *intrinsic* noise, i.e., noise resulting from non-trivial gene locus dynamics [19]. Another model describes *extrinsic* noise, i.e., noise resulting from cell-to-cell differences in rate parameters (Figure 2c). These two sources have been studied simultaneously since the early 2000s [19,20]; however, full analytical solutions have been limited to rather simple cases due to substantial mathematical complexity.

**Figure 2:**
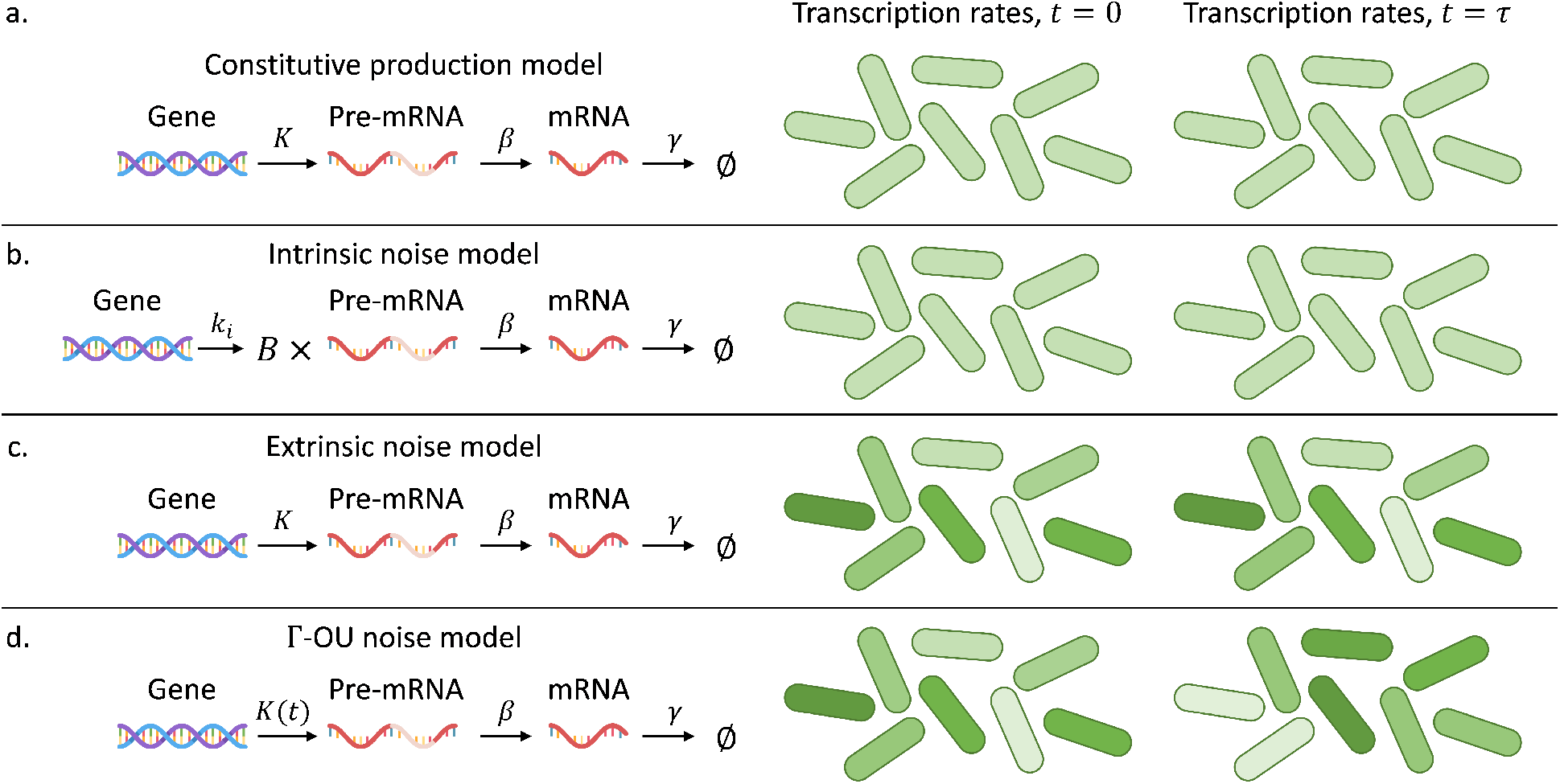
(a) Schema of the intrinsic noise model (*k*_*i*_: burst frequency; *B*: burst size drawn from a geometric distribution; *β*: pre-mRNA splicing rate; *γ*: mRNA degradation rate. Uniform shade of green indicates identical parameter values across all cells). (b) Schema of the extrinsic noise model (*K*: transcription rate; *β*: pre-mRNA splicing rate; *γ*: mRNA degradation rate. Different shades of green indicate different, but time-independent, values of *K* across cells). (c) Schema of the constitutive production model (*K*: transcription rate; *β*: pre-mRNA splicing rate; *γ*: mRNA degradation rate. Uniform shade of green indicates identical parameter values across all cells). (d) Schema of the Γ-OU noise model (*K*(*t*): transcription rate; *β*: pre-mRNA splicing rate; *γ*: mRNA degradation rate. Different shades of green indicate different values of *K* across cells and throughout time).

In the broader context of Figure 1, the more complex transcriptional models described above arise in a natural way from the simpler models by loosening assumptions and replacing constants by stochastic processes. For example, the switching model is equivalent to the birth-death process with a production rate given by the two-state telegraph process, whereas time-homogeneous extrinsic noise models arise by replacing a parameter with a static (non-changing) process initialized at a random value.

Finally, although we describe a method for the simulation of an SDE-driven CME in the current work, there is a wealth of literature using continuous models of gene expressions, essentially treating concentrations rather than single-molecule counts. A standard result in the field [21] describes an equivalence between these two approaches: given a discrete CME, a continuous Fokker-Planck equation (FPE) can be recovered via generating functions; given an FPE, the solution to a CME immediately follows via Poisson mixing. This connection is frequently exploited to make CME systems tractable [22].

### 2.2 Motivation

A recent report considers the extrinsic noise description in the discrete framework [9]. The authors use the gamma distribution to model the transcription rate distribution and motivate the choice by previous results in protein production modeling [11]. The line of reasoning posits that a set of stochastic processes induces a stationary gamma law; therefore, the stationary behavior of a CME under extrinsic noise can be simply computed as a mixture distribution. Specifically the molecule copy numbers are governed by a heterogeneous birth-death process, the stationary distribution is Poisson [7]; if the Poisson rate is, in turn, gamma-distributed, the mixed stationary distribution is negative binomial. The motivation is well in line with previous work [8], which uses the Central Limit Theorem to explain the log-normal distribution of parameter values as a natural consequence of multiplicative effects.

However, upon inspection, this description raises further questions. Although an explanatory process is invoked to motivate the choice of distribution, the process dynamics are disregarded. This is an acceptable approximation in the limit of extremely large time-scale separation – such that the process is substantially slower than the mRNA dynamics, leading to local equilibration of the processes downstream of the gene locus – but it cannot be expected to hold in all timescale regimes. Therefore, we seek to investigate the behavior of CME systems under stochastic driving at the gene locus. To exactly match the asymptotically slow extrinsic noise regime, the process *K*(*t*) describing the transcription rate must have a gamma stationary distribution; to reflect transient dynamics, it must also have nontrivial trajectories. The natural representation is a mean-reverting process described by a stochastic differential equation (SDE). Therefore, the problem requires the coupling of a CME to an SDE, which is rather nontrivial. Although analogous problems have been explored in the domain of multi-scale modeling over the past twenty years [23,24], analytical solutions are rare, and none appear to be available for this model. Even simulation is challenging: standard methods tend to use models with Brownian noise in the SDE, and require Euler–Maruyama stochastic integration combined with various rejection schema [24–26]. Intuitively, these models not amenable to *direct* (non-rejection) simulation [27] because the assumption of time-independent rates is broken, and the residence time must be computed using numerical integration.

To side-step this problem, we use the gamma Ornstein-Uhlenbeck (Γ-OU) model, which is well-known from quantitative finance [28] and has recently been applied in an investigation of intrinsic noise, albeit with rather different gene dynamics [18]. The Γ-OU model, which does not have a Brownian noise component, affords a semi-analytical solution for the state residence time, and thus enables simulation through a variant of Gillespie’s direct method [27]. We describe an algorithm to compute exact residence times, discuss several points pertaining to efficient numerical implementations, and demonstrate that the algorithm is capable of recapitulating the intrinsic, extrinsic, and constitutive models in several degenerate parameter regimes.

## 3 Notation

### 3.1 Probability distributions

The continuous uniform distribution is represented by 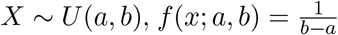, where *x ∈* [*a, b*] and *a, b ∈* ℝ. The geometric distribution is represented by *X* ∼ *Geom*(*p*), *P* (*X* = *k*; *p*) = (1 − *p*)^*k*^*p*,where *k ∈* ℕ_0_ and *p ∈* (0, 1]. The geometric distribution is well-known to arise in the short-burst limit of the two-state transcription model [2]. The negative binomial distribution is represented by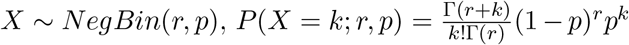, where *k ∈* N_0_, *p ∈* [0, 1], and *r >* 0. We note that MATLAB and the NumPy library use the opposite convention, with a 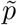 parameter defined as 1 − *p*. The exponential distribution is represented by *X* ∼ *Exp*(*η*), *f* (*x*; *η*) = *ηe*^−*ηx*^, where *x, η >* 0. This is the rate parametrization. We note that MATLAB and the NumPy library use the inverse scale parametrization with parameter *θ* = *η*^−1^. The gamma distribution is represented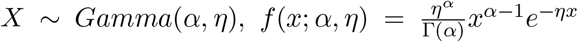, where *x, α, η >* 0. This is the shape/rate parametrization. We note that MATLAB and the NumPy library use the inverse shape/scale parametrization with parameter *θ* = *η*^−1^. In the literature, the rate *η* is usually given the variable name “*β*”; however, we use the current convention to prevent confusion with the splicing rate parameter. *Exp*(*η*) is equivalent to *Gamma*(1, *η*).

### 3.2 Stochastic processes

We follow the mathematical finance convention for the Γ-OU process [18, 29]. Specifically, a generalized OU process *K*(*t*) is the solution of the SDE

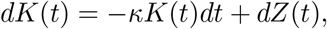

where *κ >* 0, *K*(0) = *K*_0_ *P* -almost surely, and *Z* is a subordinator of choice [30]. The Γ-OU process uses the compound Poisson subordinator 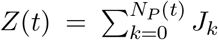, where *N*_*P*_ (*t*) is a Poisson counting process with rate *λ*, and independent random jump sizes *J*_*k*_ ∼ *Exp*(*β*). The previously reported solution [30] yields

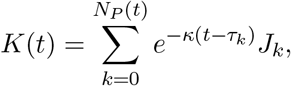

where *τ*_*k*_ are the jump times of *N*_*P*_. Note that *J*_0_ := *K*_0_ and *τ*_0_ := 0. The resulting stationary distribution is *Gamma*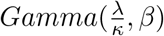.

## 4 Simulation design

### 4.1 Simulation of the Γ-OU process

We consider the standard case of simulation on *t ∈* [0, *T*]. The number of Poisson arrivals in this interval follows from the definition of a Poisson process: *N*_*P*_ (*T*) ∼ *Poisson*(*λT*). It is well-known [31] that the arrival times of a Poisson counting process on *t ∈* [0, *T*] are identically distributed to the rank statistics of a uniformly distributed random variable. Therefore, given *N*_*P*_ (*T*) total jumps, their times *τ*_*k*_, *k >* 0 can be computed by drawing *N*_*P*_ (*T*) random numbers from *U* (0, *T*) and sorting the resulting values. The jump sizes *J*_*k*_, *k >* 0 are computed by drawing *N*_*P*_ (*T*) exponential random variables with rate *η* or mean *η*^−1^. Given an initial condition, the total number of jumps, their arrival times, and their magnitudes, the Γ-OU process path is fully determined and can be easily computed.

### 4.2 Simulation of the CME

We consider a birth-death system with a single time-inhomogeneous birth rate. For illustration, we consider *n*ascent and *m*ature mRNA species, with respective instantaneous counts *n*_*n*_ and *n*_*m*_. Specifically, we consider three reactions: production with rate *a*_1_ = *K*(*t*), splicing with overall rate *a*_2_ = *βn*_*n*_, and degradation with overall rate *a*_3_ = *γn*_*m*_. Extensions to more general schema for processing downstream of transcription are trivial. The algorithm is outlined below. The full derivation, including the formula for *τ* at each step and numerical considerations for implementation, is provided in Section S2.

1. Set *t* = 0. Initialize *n*_*n*_ and *n*_*m*_.
2. Generate two uniform random variables *u*_1_ and *u*_2_.
3. Compute time step *τ* that meets the criterion 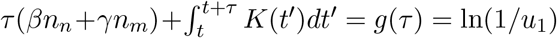.
  a. Check whether the criterion *g*(*τ*) *>* ln(1*/u*_1_) at the next jump in transcription rate:
    i. if so, use the Lambert *W* function to explicitly compute *τ*,
    ii. If not, check the next jump.
4. Compute instantaneous reaction rates *a*_*µ*_, *µ ∈ {*1, 2, 3*}*.
5. Compute net state efflux rate 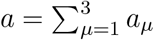
6. Select reaction index *µ* to be the lowest *i* such that 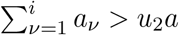
7. Advance time by *τ* .
8. Modify state variables according to the value of *µ*:
  8.1. *μ* = 1, *n*_*n*_ ← *n*_*n*_ + 1
  8.2. *μ* = 2, *n*_*n*_ ← *n*_*n*_ − 1, *n*_*m*_ ← *n*_*m*_ + 1
  8.1. *μ* = 3, *n*_*n*_ ← *n*_*n*_ − 1
9. Return to step 2.

### 5 Results

### 5.1 Asymptotic regimes

In the current section, we consider several asymptotic regimes where the SDE-driven model reduces to common physiological models of transcription. In all regimes, we simulate 10^4^ cells and use the downstream process rate parameters *β* = 1.2 and *γ* = 0.7. The stationary distributions are shown in Figure 3.

**Figure 3:**
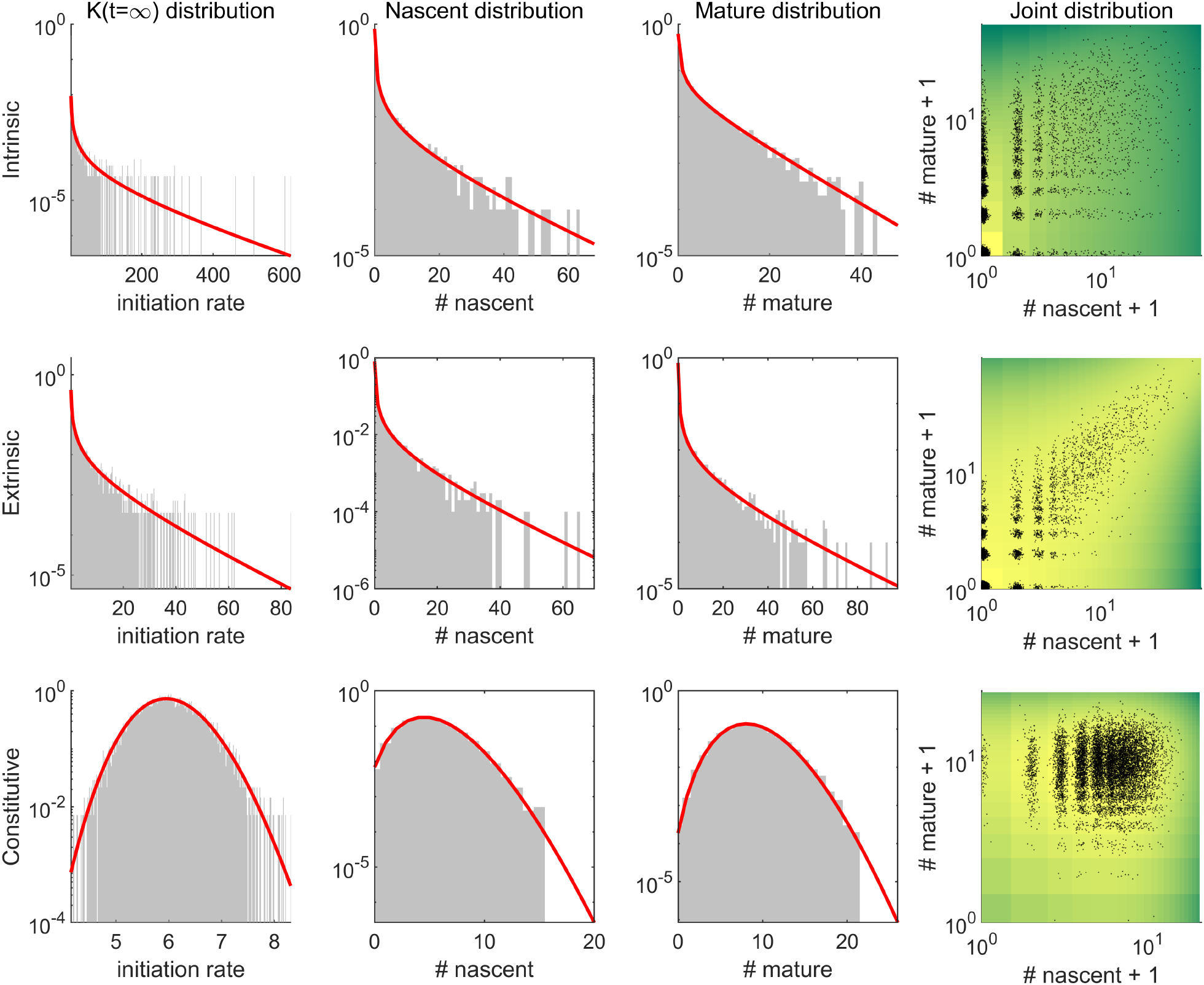
Simulation results in three asymptotic regimes (grey histograms: observed distributions; red lines: analytical results; black points: cells; right row color: log analytical joint PMF). [Code]

#### 5.1.1 Intrinsic-only noise

As *κ* → *∞* and *η* → 0, the Γ-OU stochastic process reduces to a series of peaks of infinitesimally short duration. If *κη* → *b*, a finite quantity, the mass of each peak is finite and given by *J*_*i*_*/κ*. This case reduces to the bursty system studied by Amrhein et al. [18], with burst arrival rate *λ* and mean burst size *b*. The agreement in this domain is shown in the first row of Figure 3. The time series is provided in Figure S1. The parameters used for the simulation are *κ* = 10, *λ* = 0.1, and *η* = 6.7 *×* 10^−3^, with effective mean burst size *b* = 15.

#### 5.1.2 Extrinsic-only noise

Purely extrinsic noise is conventionally modeled as a mixture with time-independent, gamma-distributed transcription parameters. In the context of transcription governed by a Γ-OU process, this corresponds, intuitively enough, to a regime with significant timescale separation between the gene locus noise and the downstream processing. Specifically, if *κ* ≪ *β, γ*, the cells experience local equilibrium. To yield a non-degenerate stationary distribution of rates, *λ* must also vanish.

Therefore, extrinsic noise is recapitulated whenever SDE dynamics are sparse and slow compared to downstream kinetics. The agreement in this domain is shown in the second row of Figure 3. The time series is provided in Figure S2. The parameters used for the simulation are *κ* = 0.12, *λ* = 0.01, and *η* = 6.7 *×* 10^−2^.

#### 5.1.3 Constitutive production

The stationary distribution rates are distributed per *Gamma*(*λ/κ, η*), with mean 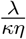and variance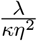. Therefore, if *η* → *∞* while *λ/κ* → *∞*, with the finite constraint 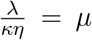, the stationary distribution reduces to a Dirac δ distribution with a point mass at *µ*. This degenerate case yields constitutive production with a constant transcription rate *µ*. The downstream nascent and mature mRNA distributions are given by *N ~ Poisson*(*µ/β*) and *M* ∼ *Poisson*(*µ/γ*), in accord with standard results [7]. The agreement in this domain is shown in the third row of Figure 3. The time series is provided in Figure S3. The parameters used for the simulation are *κ* = 8.3 *×* 10^−4^, *λ* = 0.1, and *η* = 20, with effective mean initiation rate *µ* = 6.

### 5.2 General parameter regime

The algorithm permits the exact, direct simulation of the SDE-driven system with arbitrary dynamics. The results of a sample simulation with comparable rates are shown in Figure 3. The transcriptional rate agrees with the intended theoretical form, both throughout the time series and at steady state. Furthermore, the observed long-term nascent and mature mRNA means agree with the theoretical stationary expectations 𝔼[*K*]*/β* and 𝔼[*K*]*/γ*. We do not derive expressions for the joint or marginal mRNA distributions, but note that the marginals agree fairly well with a negative binomial fit.

## 6 Discussion

We have developed an exact and direct routine to simulate an SDE coupled to a birth-death process and discussed its reduction to a set of qualitatively distinct and physiologically relevant gene expression regimes, as shown in Figure 3. Furthermore, it is suitable for simulations in intermediate parameter regimes, with a sample simulation depicted in Figure 4. In the current section, we suggest extensions to broader classes of stochastic processes, as well as theoretical directions and implications.

**Figure 4:**
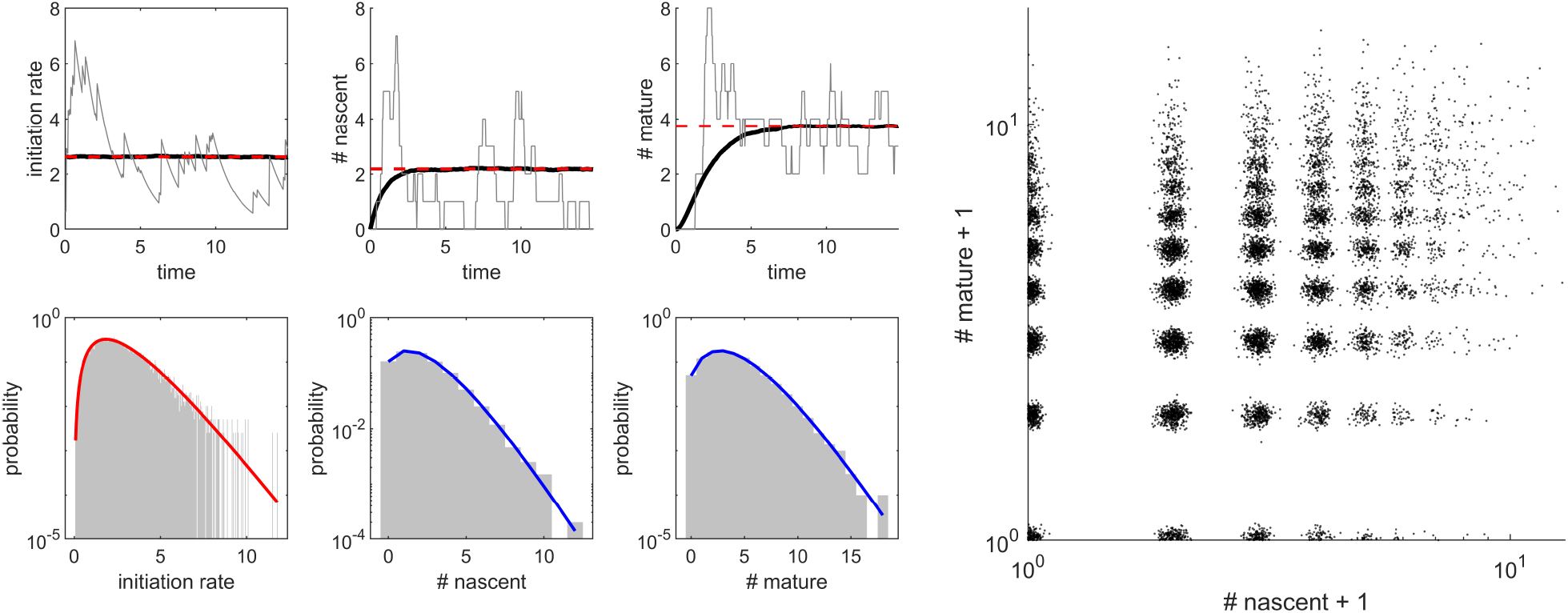
Sample simulation results (*κ* = 0.6765, *λ* = 2.3, *η* = 1.3, *β* = 1.2, *γ* = 0.7, 10^4^ cells). Top row, left: time series for initiation rate, number of nascent mRNA, number of mature mRNA (black line: mean of all iterations; grey line: single iteration, red dashed line: expected stationary mean). Bottom row, left: stationary distributions (grey histogram: observed distribution; red line: expected analytical distribution; blue line: best negative binomial fit). Right: empirical joint distribution (black points: cells; normal jitter with σ = 0.05 added).

### 6.1 Generalizations to a broader class of subordinators

Due our focus on stationary behavior under gamma-distributed transcriptional parameters, we only explore a fairly limited domain of *K*(*t*) behaviors. However, as evident from the functional form, transcriptional dynamics driven by *any* subordinator *Z*(*t*) containing a finite number of jump in every closed interval can be simulated using an identical procedure. Specifically, the core loop does not depend on the details of the random number generation process that produces *N*_*P*_ (*t*), *τ*_*k*_, and *J*_*k*_. Therefore, a fairly wide range of *K*(*t*) dynamics can be pursued with minimal modifications to the algorithm.

The case of time-varying *η* is trivial to implement. Given a function which defines *η*(*t*), it is only necessary to draw *N*_*P*_ (*T*) exponential random variables with rates 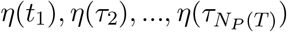. The resulting simulation is exact for analytical *η*(*t*). It may be approximate for more complex forms of *η*(*t*) given by, e.g., deterministic or stochastic differential equations. The case of time-varying *λ* is likewise straightforward; procedures for the simulation of inhomogeneous Poisson processes are readily available [32]. Parenthetically, we note that if *λ*(*t*) has an analytical integral, its simulation is equivalent to the exact Gillespie simulation of a pure-birth CME system; therefore, methods described elsewhere in the report can be used to exactly simulate the process arrival times.

The simulation is by no means limited to the simple two-step BDP illustrated here. In fact, the form of the root-finding problem in *τ* that gives rise to Equation S1 naturally suggests that the birth process can be combined with any number of time-homogeneous downstream processes. Therefore, the framework is immediately applicable to arbitrary downstream reaction graphs.

By the thinning property of a Poisson process, the simulation of a single gene locus with arrival rate *λ* is equivalent to the simultaneous simulation of *n* gene loci with rates *λ*_1_, *λ*_2_, …, *λ*_*n*_. The event– locus assignments are performed by drawing from a categorical distribution (with PMF *p*_*i*_ = *λ*_*i*_*/λ*). Finally, given different rates *η*_*i*_ at each locus, the overall compound Poisson process is produced by drawing from the distribution *Exp*(*η*_*i*_) at each event *i*.

### 6.2 Theoretical directions

Our unified description of noise sources presents an alternative to phenomenological additive or multiplicative noise models [10]. Further appeal lies in the functional form of the Γ-OU model, which permits interpretation in terms of site exposure followed by linear occlusion.

We suggest that the intermediate regime of SDE parameters, with moderate rates *κ* and *λ*, presents a natural area for exploration, as it interpolates between the intrinsic and extrinsic noise regimes. Qualitatively, the intrinsic noise regime corresponds to dispersion purely governed by the spike masses and arrivals, whereas the extrinsic noise corresponds to dispersion purely governed by spike magnitudes and the ratio of *K*(*t*) rates. We speculate that the intermediate regime corresponds to dispersion greater than either extreme allows, and thus permits the integration of both noise models in a single, self-consistent mechanistic description.

Although specific analytical solutions are outside of the scope of the current investigation, we suggest that the apparent agreement in Figure 4 (blue lines with best negative binomial fits) is only approximate, at least for the mature marginal. This hypothesis is based on the imperfect agreement between the mature marginal and the negative binomial distribution in the limit of pure intrinsic noise [33].

Throughout the current work, we focus upon the two-stage BDP. As recently discussed [16], the use of a multi-stage model yields strikingly discordant results under the assumptions of pure intrinsic and extrinsic noise. Conversely, the availability of multimodal data presents opportunities for model discrimination, even at steady state. Therefore, we suggest that multimodal information may be highly informative in intermediate regimes.

## 7 Code availability

MATLAB code that can be used to reproduce Figures 4-S3, including the simulation and plotting routines, is available at https://github.com/pachterlab/GP_2021.

## 8 Acknowledgments

The DNA, pre-mRNA, and mature mRNA illustrations used in Figure 2, reproduced from [34], are derivatives of the DNA Twemoji by Twitter, Inc., used under CC-BY 4.0. G.G. and L.P. were partially funded by NIH U19MH114830.

## S1 Supplementary figures

**Figure S1:**
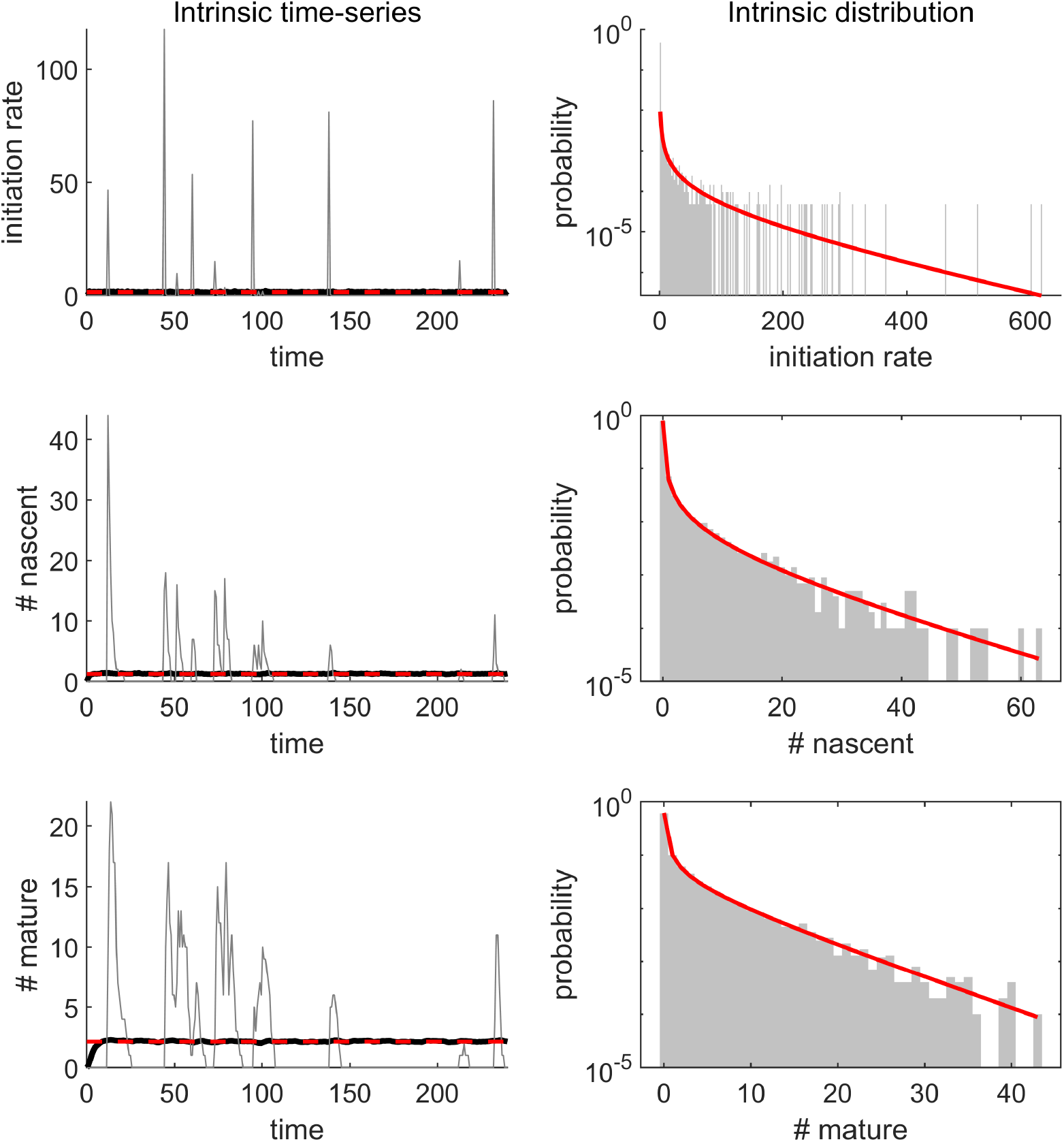
Simulation results in the intrinsic noise regime. Left: time series for initiation rate, number of nascent mRNA, number of mature mRNA (black line: mean of all iterations; grey line: single iteration, red dashed line: expected stationary mean). Right: stationary distributions (grey histogram: observed distribution; red line: expected analytical distribution).

**Figure S2:**
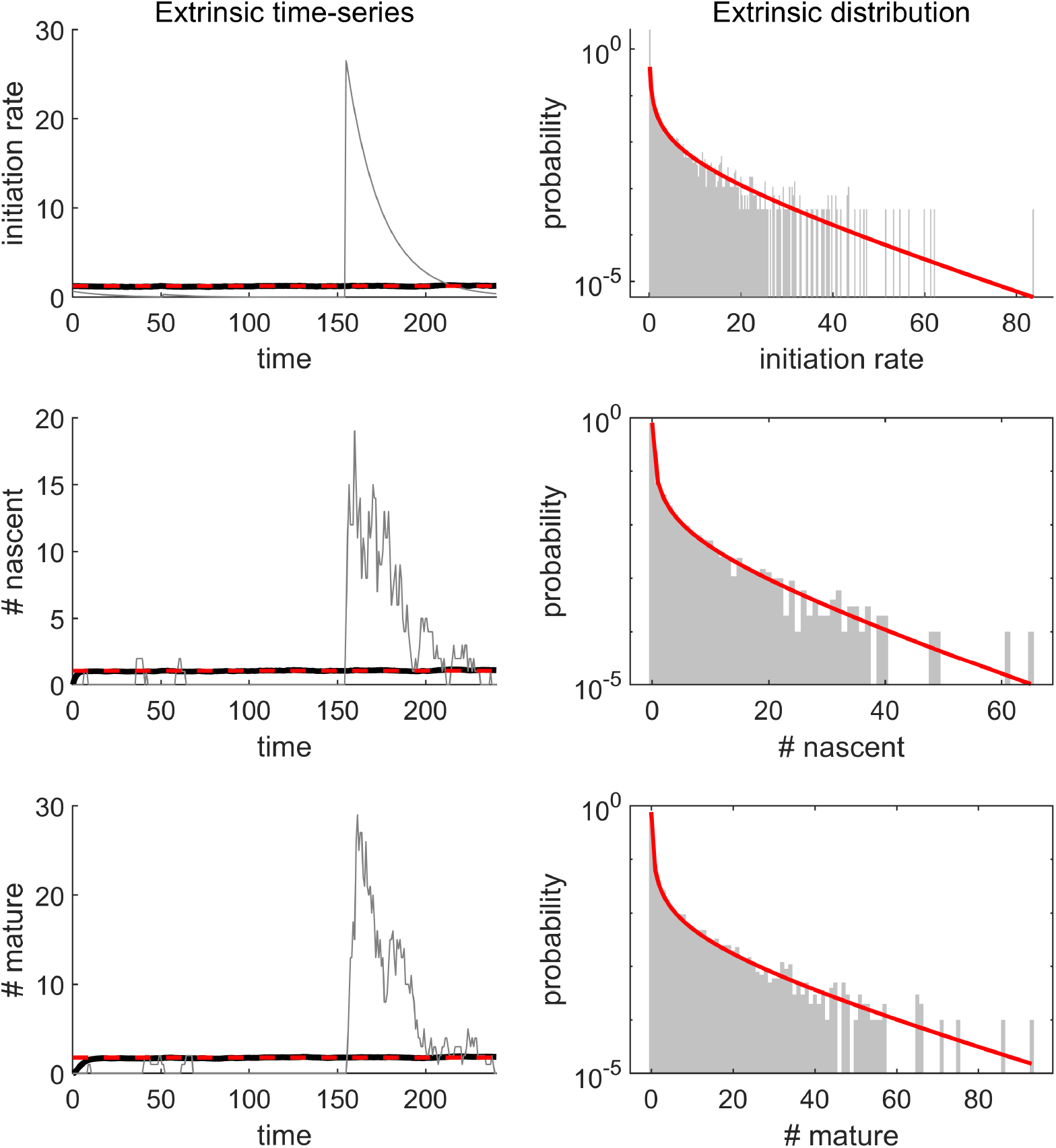
Simulation results in the extrinsic noise regime. Left: time series for initiation rate, number of nascent mRNA, number of mature mRNA (black line: mean of all iterations; grey line: single iteration, red dashed line: expected stationary mean). Right: stationary distributions (grey histogram: observed distribution; red line: expected analytical distribution).

**Figure S3:**
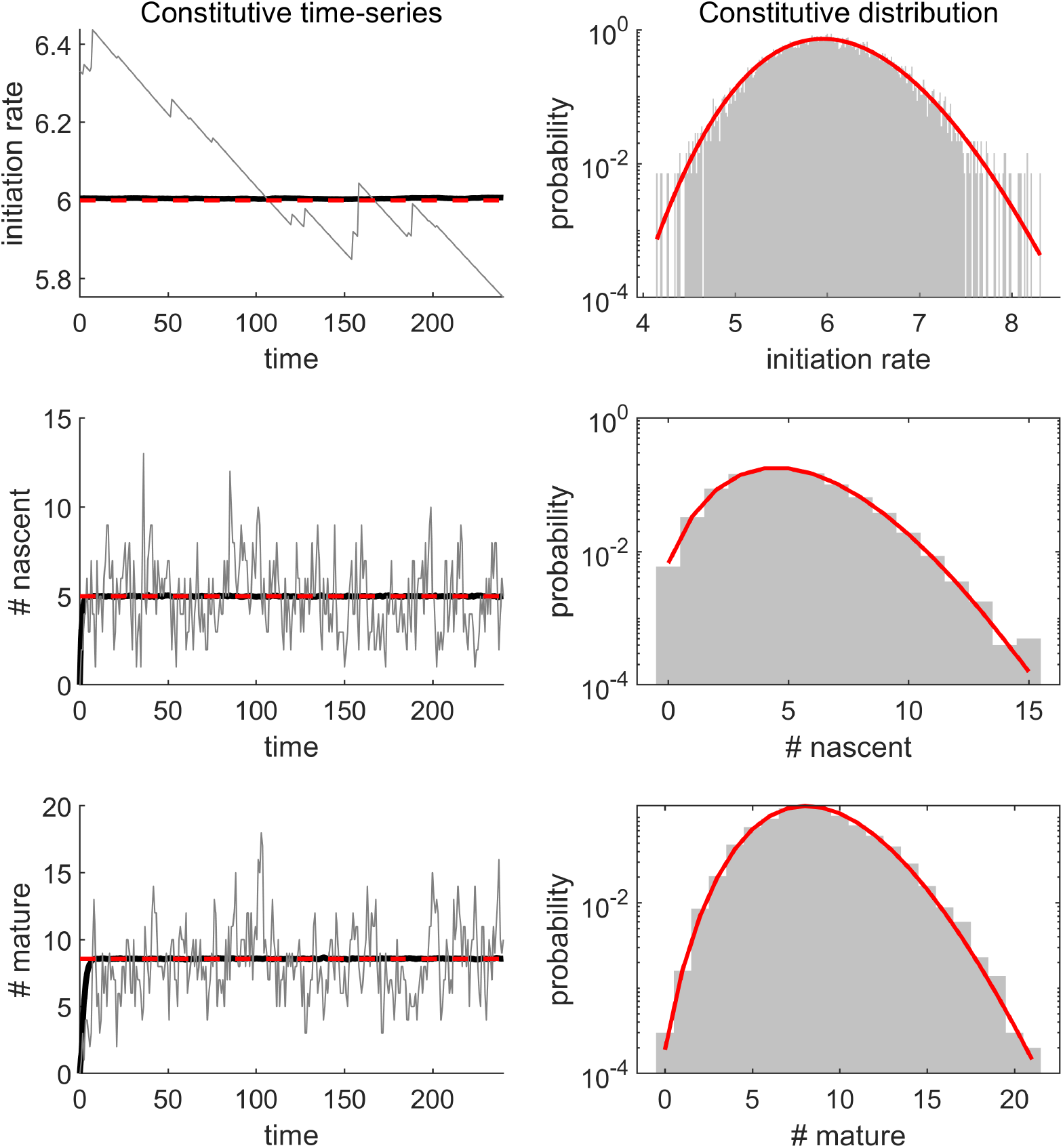
Simulation results in the constitutive production regime. Left: time series for initiation rate, number of nascent mRNA, number of mature mRNA (black line: mean of all iterations; grey line: single iteration, red dashed line: expected stationary mean). Right: stationary distributions (grey histogram: observed distribution; red line: expected analytical distribution).

## S2 Supplementary information

### S2.1 Time-homogeneous algorithm

For comparison, the standard time-homogeneous stochastic simulation algorithm (Gillespie algorithm) proceeds as follows [27]:

1. Set *t* = 0. Initialize *n*_*n*_ and *n*_*m*_.
2. Compute instantaneous reaction rates *a*_*µ*_, *µ ∈ {*1, 2, 3*}*.
3. Compute net state efflux rate 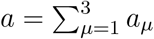
4. Generate two uniform random variables *u*_1_ and *u*_2_.
5. Compute time step 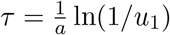
6. Select reaction index *µ* such that 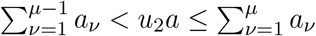
7. Advance time by *τ* .
8. Modify state variables according to the value of *µ*:
  8.1. *μ* = 1, *n*_*n*_ ← *n*_*n*_ + 1
  8.2. *μ* = 2, *n*_*n*_ ← *n*_*n*_ − 1, *n*_*m*_ ← *n*_*m*_ + 1
  8.1. *μ* = 3, *n*_*n*_ ← *n*_*n*_ − 1
9. Return to step 2.

Since the computation of *τ* presupposes constant reaction rates, this algorithm is inappropriate for the time-inhomogeneous case.

### S2.2 Time-inhomogeneous algorithm

The case of a time-inhomogeneous birth rate necessitates a more complex coupled computation [35]. Specifically, the random time step *τ* is selected according to 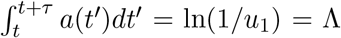. Using the definition of *a*:

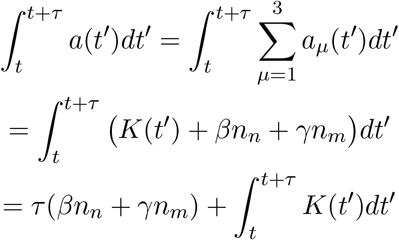

Given a particular realization, we can directly integrate *K*. Specifically:

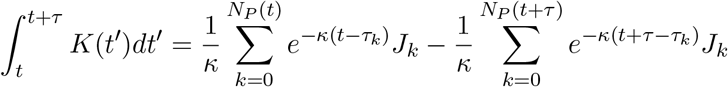

This quantity is straightforward to evaluate. However, the specific functional form makes it challenging to compute *τ* without resorting to numerical root-finding algorithms. Therefore, an alternative approach is desired for fast computation.

We begin by treating the simplest case. If *t > τ*_*k*_ for all *k*, no more jumps occur after the current time, and *K*(*t* + *τ*) exponentially decays as a function of *τ*, with the functional form *K*(*t* + *τ*) = *K*(*t*)*e*^−*κτ*^. Therefore,

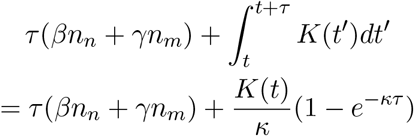

This implies the root-finding problem in *τ* :

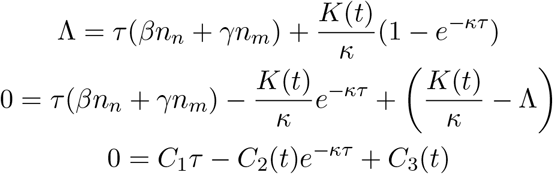

This equation has the analytical solution:

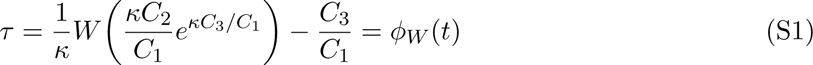

where *C*_1_, *C*_2_, and *C*_3_ are evaluated at *t*, whereas *W* is the product logarithm function, i.e. *W*_0_, the principal branch of the Lambert *W* function. This solution is straightforward to compute using standard packages, such as the MATLAB Symbolic Toolbox and the SciPy library for Python. The alternative formulation is relevant when *C*_1_ = 0:

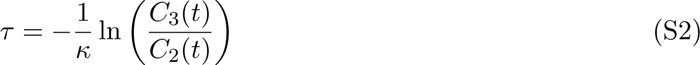

Parenthetically, we note the terminal case *t*+*τ > T*, i.e. that the reaction flux up to *T* is insufficient to match Λ. Although the SDE dynamics are not simulated past *T*, and no information about *K* is known past this time horizon, this is not a problem; the simulation remains exact up until *T*, where it halts. Another edge case, where *φ*_*W*_ (*t*) is complex-valued, implies that the total reaction flux up to *t* = *∞* is insufficient to meet Λ, and again simply leads to the termination of the simulation at*T*. This edge case only occurs when *C*_1_ = 0, as the downstream reactions occur in finite time in the converse case.

Next, we consider the first non-trivial extension: *t < τ*_*k*_ for a single *k*; a single jump occurs after the current time. For convenience of notation, we define 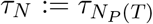. It remains to bound *t* + *τ* within the region (*t, τ*_*N*_) or the region (*τ*_*N*_, ∞).

Since 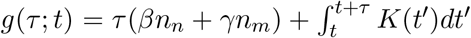is guaranteed to be monotonic, we can use a simple within the region (*t, τ*_*N*_) or the r egion (*τ*_*N*_, *∞*). binary decision procedure. If 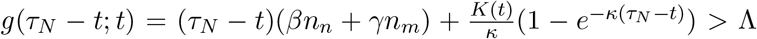, the value of the integral up to *τ*_*N*_ is an overestimate and the solution is given by Equation S1 evaluated at *t*, i.e. *φ*_*W*_ (*t*). If the converse is true, the value is an underestimate and the solution is given by *φ*_*W*_ (*τ*_*N*_) + (*τ*_*N*_ − *t*).

This procedure can be extended to an arbitrary number of jumps after *t*. The implementation requires a choice of a search procedure; we choose a simple rightward scan. Specifically, given *t < τ*_*k*_ *< τ*_*k*+1_ *<* … *< τ*_*N*_:

1. Assign upper bound for the integral *L* ← *k* and running time *t*_*R*_ ← *t*.
2. Check whether *L ≤ N* .

2.1. If *L ≤ N*, evaluate *G* = *g*(*τ*_*L*_ − *t*; *t*) = *g*(*τ*_*k*_ − *t*; *t*) + *g*(*τ*_*L*_ − *τ*_*k*_; *τ*_*k*_).

2.1.1. If *G >* Λ, the solution is given by *φ*_*W*_ (*t*_*R*_) + (*t*_*R*_ − *t*).
2.1.2. If *G <* Λ, assign *L* ← *L* + 1 and *t*_*R*_ ← *τ*_*L*_.
2.1.3. Return to 2.
2.2. If *L > N*, the solution is given by *φ*_*W*_ (*t*_*R*_) + (*t*_*R*_ − *t*).

Since *K*(*t*) is known, it is trivial to pre-compute the quantities 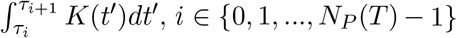 Therefore, comp uting the term *g*(*τ*_*L*_ − *τ*_*k*_; *τ*_*k*_) requires a summation over the pre-computed integral terms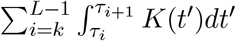 and a single evaluation of the exponential-exit product (*τ*_*L*_ − *τ*_*k*_)(*βn*_*n*_ + *γn*_*m*_). Finally, the remainder *g*(*τ*_*k*_ − *t*; *t*) requires one evaluation of the analytical integral per Gillespie time step.

With *τ* determined, it remains to select the specific reaction channel. The exponential-exit weights are given by *a*_2_ = *τβn*_*n*_ and *a*_3_ = *τγn*_*m*_. The weight *a*_1_ of the birth reaction is given by 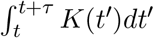which is given by

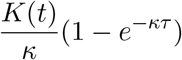

if no jumps occur up within (*t, t* + *τ*), and

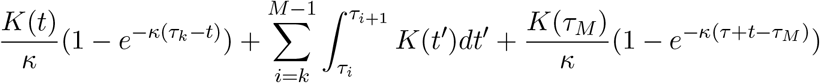

if *t < τ*_*k*_ *< τ*_*k*+1_ *<* … *< τ*_*M*_ *< t* + *τ*. Therefore, the following steps of the time-inhomogeneous algorithm are identical to steps 6-9 of the time-homogeneous algorithm.

### S2.3 Implementation details

Several points regarding the efficient implementation of the algorithm bear further discussion. For computational facility, at each step of the Gillespie simulation, we set *τ*_*k*−1_ → *t* and *K*(*τ*_*k*−1_) → *K*(*t*). This approach creates a virtual jump at the current time, and allows treating the integral 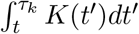 without creating a special edge case. Furthermore, to minimize the number of times the pre-computed integrals are accessed, we compute ∆*G* at each step, compare it to Λ, and *decrement*Λ by ∆*G* if the reaction flux is insufficient.

The formulation in Equation S1 is susceptible to overflow as *κC*_3_*/C*_1_ → *∞*. A naïve computation at sufficiently high values yields 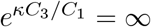 and *τ* = *∞*, halting the simulation. Therefore, wherever overflow is likely to occur, it is necessary to use the appropriate approximation to *W*. We follow the approach of Iacono and Boyd [36].

As *x* → *∞*, ln(1 + *x*) has the Puiseux series representation ln(*x*) + *x*^−1^ + *O*(*x*^−2^). For *x* sufficiently high to produce overflow, we truncate at the first term and use ln(1 + *x*) ≈ ln(*x*).

As an initial guess, we can choose *W*_0_(*x*) = ln(1 + *x*ζ(*x*)), where 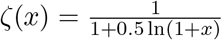; we note thatthe subscript refers to the approximation order rather than the branch of the function. Using the Puiseux series, 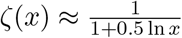. Assuming *x* is high enough, we can further assume ln(1 + *x*ζ(*x*)) ≈ln(*x*ζ(*x*)) = ln *x* − ln(1 + 0.5 ln(*x*)). Higher-order approximations follow from the iterative schema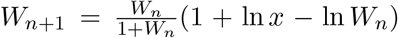. We use the =fifth-order iterative approximation whenever the argument of the Lambert *W* function is greater than 10^3^.

